# Genetic factors outside the metabolic cluster for plastid-derived sesquiterpenes are required to pursue arthropod-resistant tomatoes

**DOI:** 10.1101/2020.02.21.960112

**Authors:** Rodrigo Therezan, Ruy Kortbeek, Eloisa Vendemiatti, Saioa Legarrea, Severino M. de Alencar, Robert Schuurink, Petra Bleeker, Lázaro E. P. Peres

**Affiliations:** Laboratory of Plant Developmental Genetics. Department of Biological Sciences, “Luiz de Queiroz” College of Agriculture, University of Sao Paulo, 13418-900, Piracicaba, SP, Brazil; Green Life Science Research Cluster, Swammerdam Institute for Life Sciences, University of Amsterdam, Science Park 904, 1098 XH Amsterdam, The Netherlands; Molecular & Chemical Ecology, Institute for Biodiversity and Ecosystem Dynamics, University of Amsterdam, PO Box 94240,1090 GE Amsterdam, The Netherlands; Department of Agri-Food Industry, Food and Nutrition, “Luiz de Queiroz” College of Agriculture, University of Sao Paulo, 13418-900, Piracicaba, SP, Brazil

**Keywords:** Type-VI trichomes, tomato trichomes, santalene, bergamotene, *Solanum habrochaites*, glandular trichome

## Abstract

To deal with arthropod pests the tomato wild relatives produce a variety of defense compounds in their glandular trichomes. In *Solanum habrochaites* LA1777, a functional cluster of genes on chromosome 8 controls plastid-derived sesquiterpene synthesis not found in cultivated tomatoes. The main genes at the cluster are *Z-prenyltransferase (zFPS)* that produces *Z-Z-*farnesyl diphosphate (*Z,Z*-FPP), and *Santalene and Bergamotene Synthase* (*SBS*) that uses *Z,Z-*FPP to produce α-santalene, β-bergamotene, and α-bergamotene in type-VI glandular trichomes. Both LA1777 and cultivated tomatoes have type-VI trichomes, but the gland in cultivated tomato is much smaller containing low levels of monoterpenes and cytosolic-derived sesquiterpenes, which do not provide tomato with the same pest resistance as in LA1777. We successfully transferred the plastid-derived sesquiterpene pathway from LA1777 to type-VI trichomes of a cultivated tomato (cv. Micro-Tom, MT) by a back-crossing approach. The trichomes of the introgressed line named MT-*Sesquiterpene synthase 2* (MT-*Sst2*) produced even higher levels of α-santalene, β-bergamotene, and α-bergamotene than the type-VI glandular trichomes of LA1777. We also noticed that the type-VI trichome internal storage-cavity size increases in MT-*Sst2*, probably as an “inflated balloon” effect of the increased amount of sesquiterpenes. Surprisingly, the presence of high amounts of plastid-derived sesquiterpenes was not sufficient to confer resistance to various tomato pests in MT-*Sst2*. Since MT-*Sst2* made the same sesquiterpenes as LA1777, this points to additional factors, outside the genomic region thought to be the metabolic cluster, necessary to obtain arthropod-resistant tomatoes. Our results also provide a better understanding of the morphology of *S. habrochaites* type-VI trichomes.

**One-sentence summary:** Cultivated tomatoes harboring the plastid-derived sesquiterpenes from *S. habrochaites* need additional genetic components necessary to convert them into effective insecticides.

## INTRODUCTION

Isoprenoids are the most abundant and diverse class of compounds produced by plants with a wide variety of biological functions (Dudareva *et al.*, 2013). They are produced through combining multiple five-carbon units (C5) of isoprene and are essential for plant growth and development. They participate as precursors for several components of essential processes like photosynthesis, respiration, cell cycle control (Estévez *et al.*, 2001) and plant hormones such as gibberellins, abscisic acid, brassinosteroids and strigolactones (Falara *et al.*, 2011). Isoprenoids also play an important role in the interactions of plants with the environment, including defense against herbivorous insects and attraction of pollinators (Dudareva *et al.*, 2013).

In plants, all isoprenoids originate from two distinct metabolic pathways: the mevalonate (MVA) pathway, located in the cytosol and the 2-C-methyl-D-erythritol 4-phosphate (MEP) pathway located in plastids. Both pathways produce isopentenyl diphosphate (IPP) and dimethylallyl diphosphate (DMAPP), the isoprene building blocks used by terpene synthases (TPSs) to catalyze the formation of C10 monoterpenes, C15 sesquiterpenes or C20 diterpenes (Tholl, 2006).

In cultivated tomato (*Solanum lycopersicum*), sesquiterpene biosynthesis usually takes place in the cytosol from the MVA pathway. However, in glandular trichomes of the wild tomato species *S. habrochaites,* sesquiterpenes are also produced in the plastids from the MEP pathway (Sallaud *et al.*, 2009). The presence of plastid-derived sesquiterpenes in some wild species has been described to be responsible for the decreased damage by insects, making these wild species naturally resistant to multiple pests such as lepidopterans (Eigenbrode *et al.*, 1994; Eigenbrode *et al.*, 1996), whiteflies (*Bemisia spp.*) (Bleeker *et al.*, 2012) and also spider mites (Maluf *et al.*, 2001).

Two independent loci have been associated with the biosynthesis of two different classes of sesquiterpenes in *S. habrochaites*. The *Sesquiterpene synthase 1* (*SsT1*) locus on chromosome 6 is responsible for the accumulation of cytosol-derived sesquiterpenes. At this locus, *TPS12* is associated with β-caryophyllene and α-humulene biosynthesis and *TPS9* with the production of germacrenes. The *S. habrochaites TPS9* allele (*ShTPS9*) is associated with germacrene B and D production (Hoeven van der *et al.*, 2000; Bleeker, Spyropoulou, *et al.*, 2011; Falara *et al.*, 2011). The existence of *SlTPS9*, that makes germacrene C, was also reported for the cultivar VFNT Cherry (Colby *et al.*, 1998), but it is worth noting that this cultivar has introgressions of the wild species *S. peruvianum* on chromosome 6. The second locus harboring sesquiterpenes synthases is the *Sesquiterpene synthase 2* (*SsT2*) locus on chromosome 8. At this locus *S. habrochaites* has a cluster of *TPSs* (*TPS18, TPS20* and *TPS45*) encoding enzymes responsible for the accumulation of plastid-derived sesquiterpenes, including α-santalene, α-bergamotene, β-bergamotene and 7-epizingiberene (Sallaud *et al.*, 2009; Bleeker, Diergaarde, *et al.*, 2011). In cultivated tomato, a cluster of five functional *TPS* genes (*TPS18*, *TPS19*, *TPS20*, *TPS21*, and *TPS41*) is present in the equivalent locus on chromosome 8 (Falara et al., 2011). In addition, this same chromosomal region also contains the *Neryl Diphosphate Synthase 1* (*SlNDPS1*) gene, which codes for an enzyme catalyzing the formation of neryl diphosphate (NPP). NPP is used by tomato *TPS20* to synthesize β-phellandrene and several other monoterpenes in the plastids (Falara *et al.*, 2011; Matsuba *et al.*, 2013). In *S. habrochaites* the *cis-Farnesyl Diphosphate Synthase* (*zFPS*) gene is homologous to the *SlNDPS1* gene (Matsuba *et al.*, 2013). The *zFPS* codes for a Z-prenyltransferase that catalyzes the synthesis of *Z-Z-*farnesyl diphosphate (*Z,Z*-FPP) from IPP and DMAPP. The *TPS45* gene from *S. habrochaites* LA1777 encodes a *Santalene and Bergamotene Synthase (SBS)* that uses *Z,Z-*FPP as a substrate to produce plastid-derived sesquiterpenes (Sallaud *et al.*, 2009; Matsuba *et al.*, 2013). Both *zFPS* and *SBS* contain putative chloroplast targeting sequences allowing the biosynthesis of the sesquiterpenes in this organelle.

The *zFPS* and *SBS* genes are specifically expressed in type-VI glandular trichomes (Sallaud *et al.*, 2009) that harbor chloroplasts and are present on several tomato wild-relatives (Kang *et al.*, 2010; Glas *et al.*, 2012; Balcke *et al.*, 2017). In cultivated tomato, type-VI trichomes contain a single basal cell connected to a short (~0.1 mm) unicellular stalk which is connected by an intermediate cell to the four-celled glandular head containing chloroplasts and other organelles (Bergau *et al.*, 2015). In *S. habrochaites*, the stalk is longer (~0.2 mm) and the trichome has a round glandular head instead of visible 4 distinct cells (Besser *et al.*, 2009).

In general, cultivated tomatoes are highly vulnerable to several arthropod pests, which include whiteflies, spider mites, and thrips. In addition, some pest, such as whiteflies can spread viruses (Moodley *et al.*, 2019). Under heavy infestation, these pests can cause a reduction of plant vigor and yield which can lead to huge losses in productivity (Wakil *et al.*, 2018). Consequently, to minimize the damage caused by pests, high amounts of pesticides have been applied in agri- and horticulture (Silva *et al.*, 2011). In this sense, an alternative to chemical pest control could be the use of commercial tomatoes carrying favorable genetic factors from tomato wild species. Herein, we investigated whether the introduction of the genetic pathway for plastid-derived sesquiterpenes from a wild species into cultivated tomato could increase resistance to arthropod tomato pests. We show that the *Sst2* gene cluster that controls santalene and bergamotene production in *S. habrochaites* LA1777 can be effectively transferred to cultivated tomato (cv. Micro-Tom) and function in its type-VI trichomes. Tomato type-VI trichomes that accumulated high levels of plastid-derived sesquiterpenes also increased the size of the internal gland cavity, providing a better understanding of differential trichome morphology. We further demonstrated that, the high production of “wild tomato sesquiterpenes” was not sufficient to confer resistance to tomato-pests. Apparently genetic factors outside the metabolic cluster on chromosome 8 are required for the production of the anti-insect compounds in *S. habrochaites*.

## RESULTS

### Introgression of *SsT2* gene cluster from *S. habrochaites* LA1777 into *S. lycopersicum* cv. Micro-Tom (MT)

In order to introduce the plastid-derived sesquiterpene pathway into the Micro-Tom (MT) cultivar, we crossed *S. habrochaites* LA1777 with a MT line harbouring the *lutescent 1* mutation (MT-*l*1) and used it as a recurrent parent (Fig.1A). Both the *lutescent 1* mutation and the *SsT2* locus map at the short arm of the chromosome 8 (Tanksley *et al.*, 1992; Sallaud *et al.*, 2009). The *lutescent 1* phenotype comprises a premature and progressive yellowing of the leaves due to impaired chlorophyll accumulation (starting from the base of the plant) (Fig S1 A and B), a lack of chlorophyll accumulation in the pistils (Fig S1 C) and whitish-yellow fruits (Barry *et al.*, 2012). This allowed us to introgress the *S. habrochaites SsT2* locus into MT by a relatively easy visual selection of the progeny. In each generation of introgression, we selected for plants not presenting the *lutescent 1* phenotype, which implies the presence of the equivalent chromosome segment of *S. habrochaites* containing the *SsT2* locus. In the F_2_ generation, plants not harbouring the *lutescent 1* mutation were used for back-crossing (BC) with MT-*l1*. This procedure was repeated in the BC_1_ and subsequent generations. After six BC generations and self-pollination (BC_6_F_2_) using the visual marker, we employed CAPS markers based on single nucleotide polymorphisms (SNPs) for a genetic screen for homozygous plants harbouring the *SsT2* locus from the wild species. In the BC_6_F_3_ generation and generations thereafter (BC_6_F_n_), the obtained plants were considered a near isogenic line (NIL), no longer segregating for the presence of the wild sesquiterpenes pathway and other traits. The NIL was named MT-*Sesquiterpene synthase 2* (MT-*Sst2*).

**Fig. 1.**
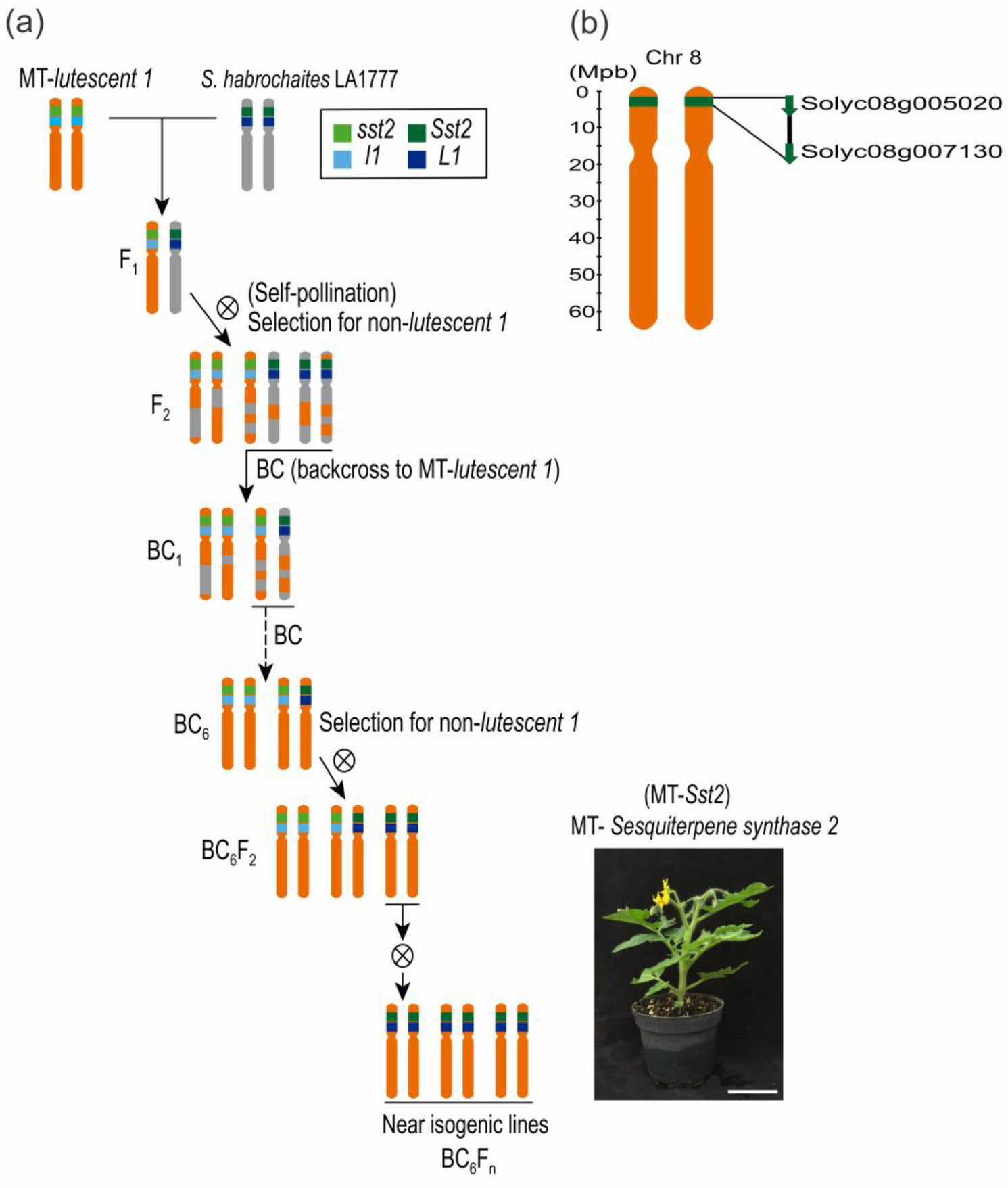
Scheme of crossing and backcrossing (BC) to create a Micro-Tom (MT) near isogenic line (NIL) harboring the *Solanum habrochaites* LA1777 genes for the “*Sesquiterpene Synthase 2*” (*SsT2*) locus. **(a)** MT NIL bearing the *lutescent 1* mutation was used to assist the introgression process as a morphological marker (the absence of the *lutescent 1* phenotype was used as an indicator of the presence of the LA1777 genes in the *SsT2* locus). The presence of MT (*l1* and *sst2*) or LA1777 (*L1* and *Sst2*) variants is indicated in different colors. **(b)** The ID (Solyc) of the genetic markers used to determine the introgression borders are depicted.

We employed CAPS markers to determine the size of the fragment introgressed into the MT background. Genetic mapping positioned the introgressed region between *Solyc08g005020* and *Solyc08g007130* on the top of chromosome 8 (Fig. 1B). The introgressed region overlaps with the mapping position of *SsT2* previously reported by Sallaud *et al.* (2009), which also confirmed that both *zFPS* and *SBS* genes were mapped to this region. The introgressed region in MT-*Sst2* is furthermore consistent with the *SsT2* locus found in a set of ILs between *S. lycopersicum* × *S. habrochaites* LA1777 that were able to produce plastid-derived sesquiterpenes (Hoeven van der *et al.*, 2000).

Additionally, we analysed the relative transcript levels of *zFPS* and *SBS* by quantitative RT-PCR in trichomes of MT-*Sst2* and *S. habrochaites* LA1777. Both genes were indeed expressed in MT-*SsT2* but transcripts levels were significantly lower in MT-*Sst2* compared to the wild parent (Fig. 2).

**Fig. 2.**
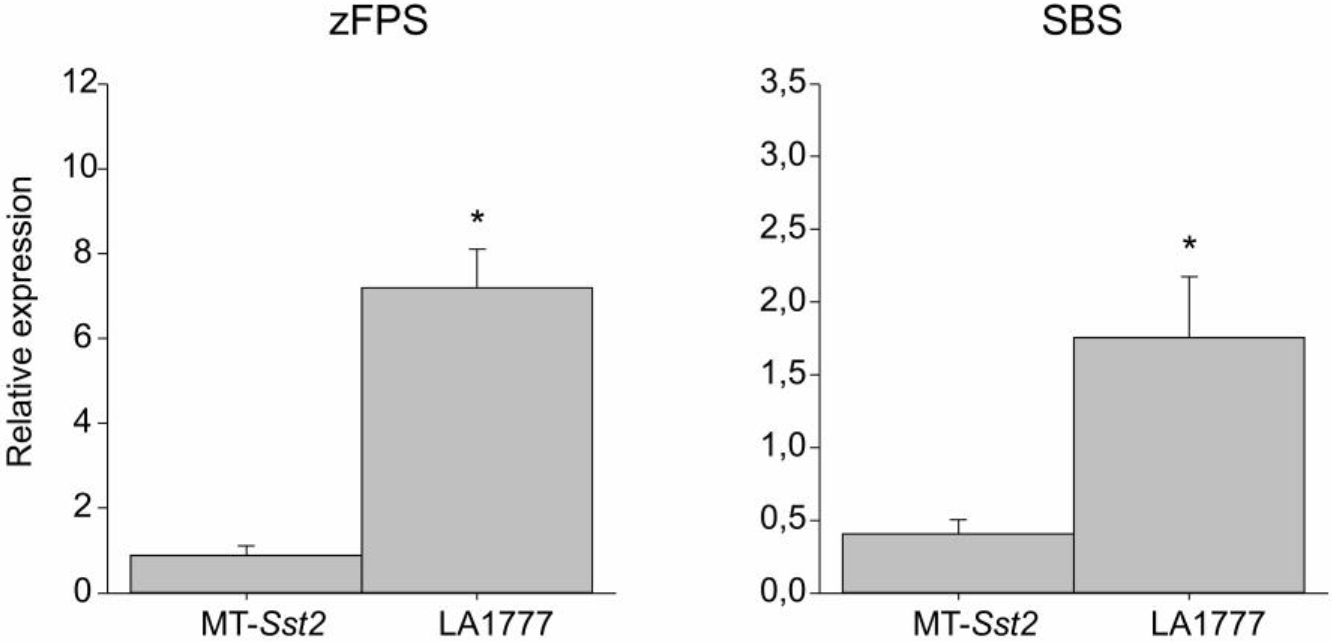
Transcript levels of the *SsT2* locus-derived genes *cis-Farnesyl Diphosphate Synthase* (*zFPS*) and *Santalene and Bergamotene Synthase* (*SBS*) in trichomes from MT-*Sst2* and *S. habrochaites* LA1777. Mean values of 4 biological replicates are shown. Transcript levels were normalized for *Rubisco conjugating enzyme 1* (RCE1). Asterisks indicate mean significantly different from MT-*Sst2*, according to Student t-test (P≤ 0.05).

### Production of the sesquiterpenes santalene and bergamotene in the Micro-Tom line harboring the *S. habrochaites SsT2* locus

To confirm the presence of the *S. habrochaites* sesquiterpene pathway in the MT-*Sst2* line, we performed Gas-chromatography Mass-Spectrometry (GC-MS) analyses on isolated type-VI trichome glands. The type-VI trichomes of the MT-*Sst2* line not only accumulated sesquiterpenes, but also produced higher amounts of these plastid-derived sesquiterpenes, compared to parental *S. habrochaites* LA1777 (Fig. 3).

**Fig. 3.**
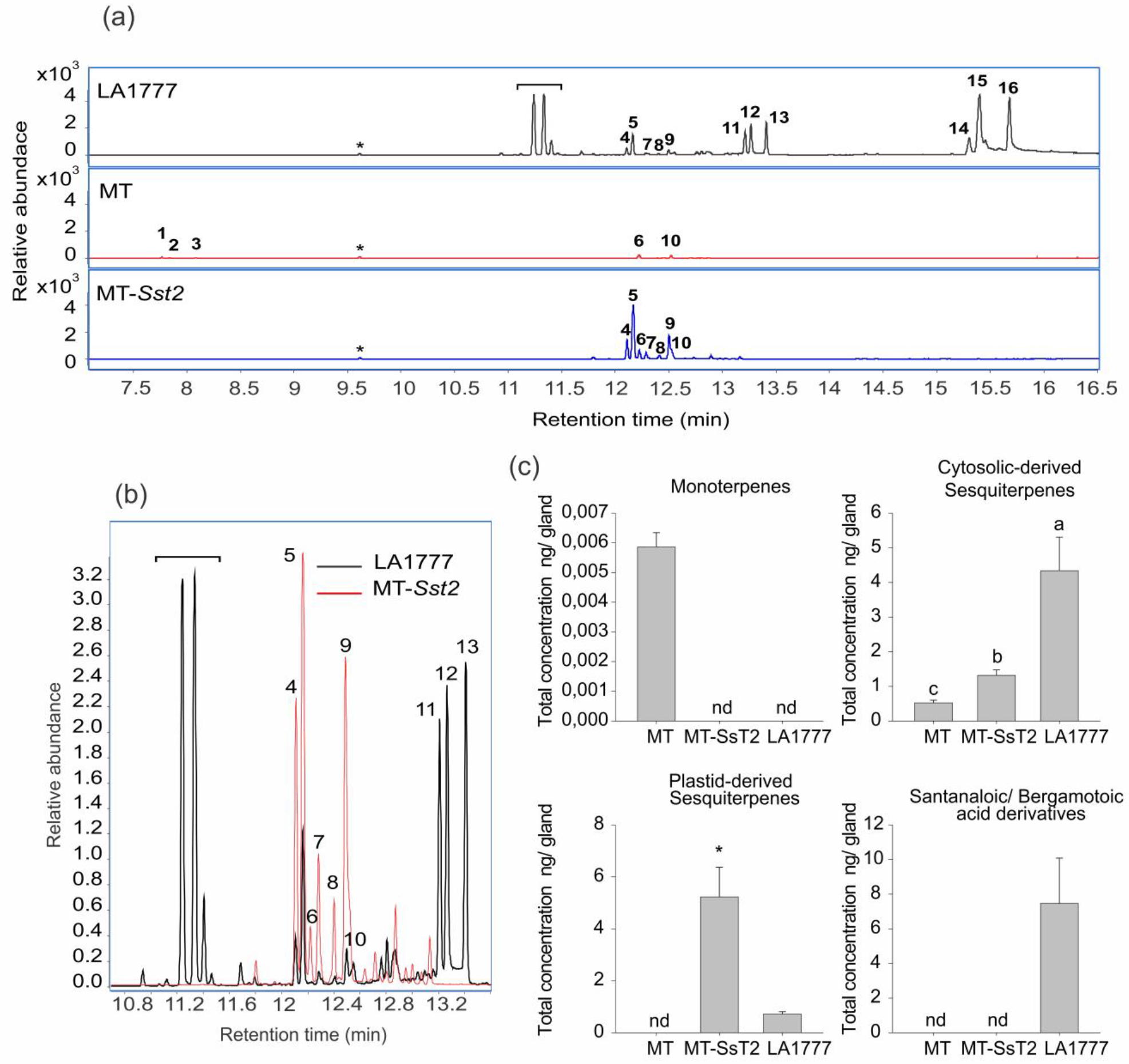
**(a)** GC-MS chromatograms showing mono and sesquiterpenes found in type-VI trichomes from Micro-Tom (MT), MT-*Sst2,* and *S. habrochaites* LA1777. The indicated peaks corresponds to the following compounds: (1) 2-carene, (2) α-phellandrene, (3) β-phellandrene/ D-limonene, (4) α-bergamotene, (5) α-santalene, (6) β-caryophyllene, (7) exo-α-bergamotene, (8) epi-β-santalene, (9) endo-β-bergamotene, (10) α-humulene, (11) germacrene B, (12) selinene, (13) germacrene D, (14) α-bergamotoic acid, (15) α-santanaloic acid and (16) β-bergamotoic acid. The asterisks indicate the peaks related to the internal standard. The bracket indicates peaks related to unidentified putatively lipid-originating compounds. The chromatogram shows the detector response for the ion mass 93.069 and 108.056. **(b)** Gas chromatograms overlaying sesquiterpenes found in type-VI trichomes from MT-*Sst2* and *Solanum habrochaites* LA1777. The bracket indicates peaks related to unidentified putatively lipid-originating compounds. **(c)** Total amount of compounds present in type-VI glandular trichomes of each genotype. Total concentration of monoterpenes: (1) 2-carene, (2) α-phellandrene and (3) β-phellandrene/ D-limonene; Total concentration of cytosolic sesquiterpenes: (6) β-caryophyllene, (10) α-humulene, (11) germacrene B, (12) selinene and (13) germacrene D; Total concentration of plastid-derived sesquiterpenes: (4) α-bergamotene, (5) α-santalene, (7) exo-α-bergamotene, (8) epi-β-santalene and (9) endo-β-bergamotene; Total concentration of santalanoic/ bergamotoic acid derivative: (14) α-bergamotoic acid, (15) α-santanaloic acid (16) β-bergamotoic. The bars represent the mean ± SE of five biological replicates. For each sample, 300 type-VI glandular trichomes were collected with a glass capillary for GC-MS analysis. Bars indicated with an asterisk were significantly different according to t-test (P≤ 0.05). Bars indicated with different letters were significantly different according to Fisher’s LSD test (P≤ 0.05) after ANOVA. nd, Not detected.

As expected, we did not detect the monoterpenes found in the MT parental in the MT-*Sst2* line (Fig. 3, (peaks 1, 2 and 3), Fig S2). Since the monoterpenes 2-carene, α-phellandrene and β-phellandrene and D-limonene are produced by enzymes encoded by *SlTPS20* genes on tomato chromosome 8, it was predicted that these monoterpenes on homozygous MT-*Sst2* plants would be replaced by the sesquiterpenes produced by enzymes encoded by the wild species alleles on the same chromosomal region. Indeed, the MT-*Sst2* line produced the plastid-derived sesquiterpenes (α-bergamotene, α-santalene, exo-α-bergamotene, epi-β-santalene, and endo-β-bergamotene), also present in *S. habrochaites* LA1777 (Fig. 3A and B). Although we found high levels of plastid-derived sesquiterpenes in the MT-*Sst2* line, we did not detect any carboxylic acid derivatives from santalene or bergamotene, as in the wild species (Fig. 3A, peaks 14, 15 and 16, and 3C). As expected, no santalene or bergamotene was present in control MT trichomes (Fig. 3A, C and S2). However, not only was there a 7-fold increase in the plastid-derived sesquiterpenes of individual type-VI trichomes of MT-*Sst2* compared to LA1777, but it appeared that the MT-*Sst2* type-VI trichomes also produce substantially (2-fold) more cytosolic-sesquiterpenes when compared to MT (Fig. 3C).

### Production of mono and sesquiterpenes in type-VI trichomes under different allelic dosages at the *SsT2* locus

In order to investigate how the allelic dosage at the *SsT2* locus affects the levels of mono and sesquiterpenes in type-VI trichomes, we selected homozygous *sst2*/*sst2* and *Sst2*/*Sst2*, and heterozygous *sst2*/*Sst2* MT plants. The molecular markers used for selection were designed based on polymorphisms found in the genomic sequence of cultivated tomato and LA1777 (Table S1). Note that there is no complete synteny between the *SsT2* locus of *S. lycopersicum*, and *S. habrochaites.* Hence, at this locus, *S. lycopersicum* contains the functional genes *TPS18*, *TPS19*, *TPS20*, *TPS21*, *TPS41* and *SlNDPS1*, whereas *S. habrochaites* contains the functional genes *TPS18*, *TPS20*, *TPS45* (*SBS*) and *zFPS* (Matsuba *et al.*, 2013). Thus, the *sst2/Sst2* plants can be better considered hemizygous for both set of genes from each parental.

The amount of the monoterpenes did not differ significantly between homozygous *sst2*/*sst2* and hemizygous *sst2*/*Sst2* plants. As predicted, monoterpenes were absent in homozygous *Sst2/Sst2* plants (Fig.4) since *SlNDPS1* and *SlTPS20* are not present in *Sst2/Sst2* plants. On the other hand, there was no significant effect of gene dosage for *SlNDPS1* when comparing monoterpene content of homozygous *sst2/sst2* and hemizygous *sst2/Sst2* plants. Regarding the concentration of cytosolic-derived sesquiterpenes β-caryophyllene and α-humulene, the hemizygous *sst2/Sst2* plants did not differ significantly from homozygous *sst2/sst2* plants, while the homozygous *Sst2/Sst2* did, produced higher amounts of these compounds, especially α-humulene (Fig.4).

**Fig. 4.**
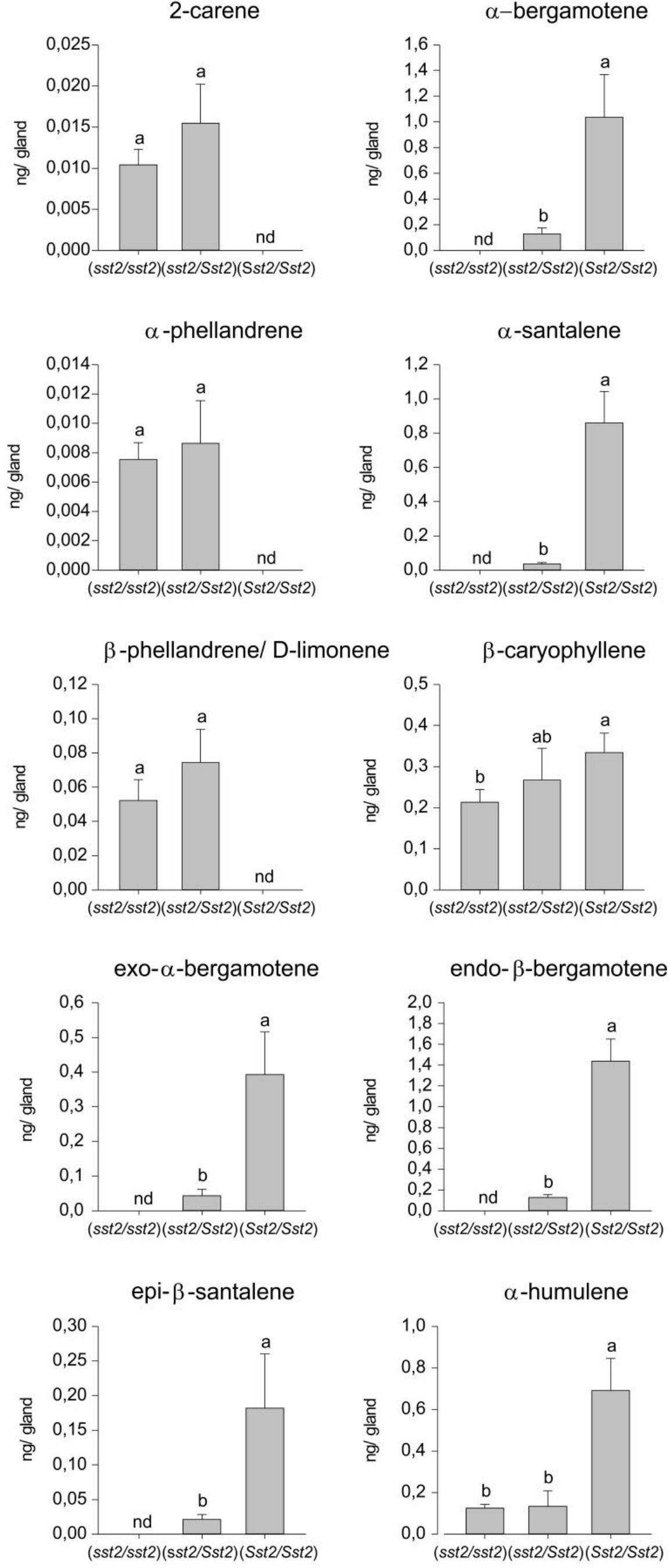
Volatile terpene levels in type-VI glandular trichomes from Micro-Tom plants homozygous (*sst2/sst2*, *Sst2/Sst2*) and hemizygous (*sst2/Sst2)* at the *SsT2* locus. The data show the amount of each compound present in type-VI glandular trichomes. Each data point represents the mean and SE of five biological replicates. For each sample, 300 type-VI glandular trichomes were collected with a glass capillary before GC-MS. Bars indicated with different letters were significantly different according to Fisher’s LSD test (P≤ 0.05) after ANOVA. nd, Not detected.

The concentrations of all plastid-derived sesquiterpenes α-bergamotene, α-santalene, exo-α-bergamotene, epi-β-santalene, and endo-β-bergamotene were higher in the homozygous *Sst2*/*Sst2* type-VI trichomes when compared to the trichomes of hemizygous *sst2*/*Sst2* plants, indicating that these compounds are under the effect of gene dosage.

### Trichome abundance, morphology and gene expression in the Micro-Tom line harboring the *S. habrochaites* genes in the *SsT2* locus

We next verified if the locus substitution caused changes in abundance of different trichome types and the morphology of type-VI trichomes in adult leaves of the MT-*Sst2* line. The densities of type-VI trichomes were not altered in MT-*Sst2* compared to MT for both adaxial or abaxial leaf surfaces (Fig. 5A and 5B). The wild species showed significantly higher numbers of type-VI trichome on the adaxial and fewer numbers on the abaxial leaf surface (Fig. 5B).

**Fig. 5.**
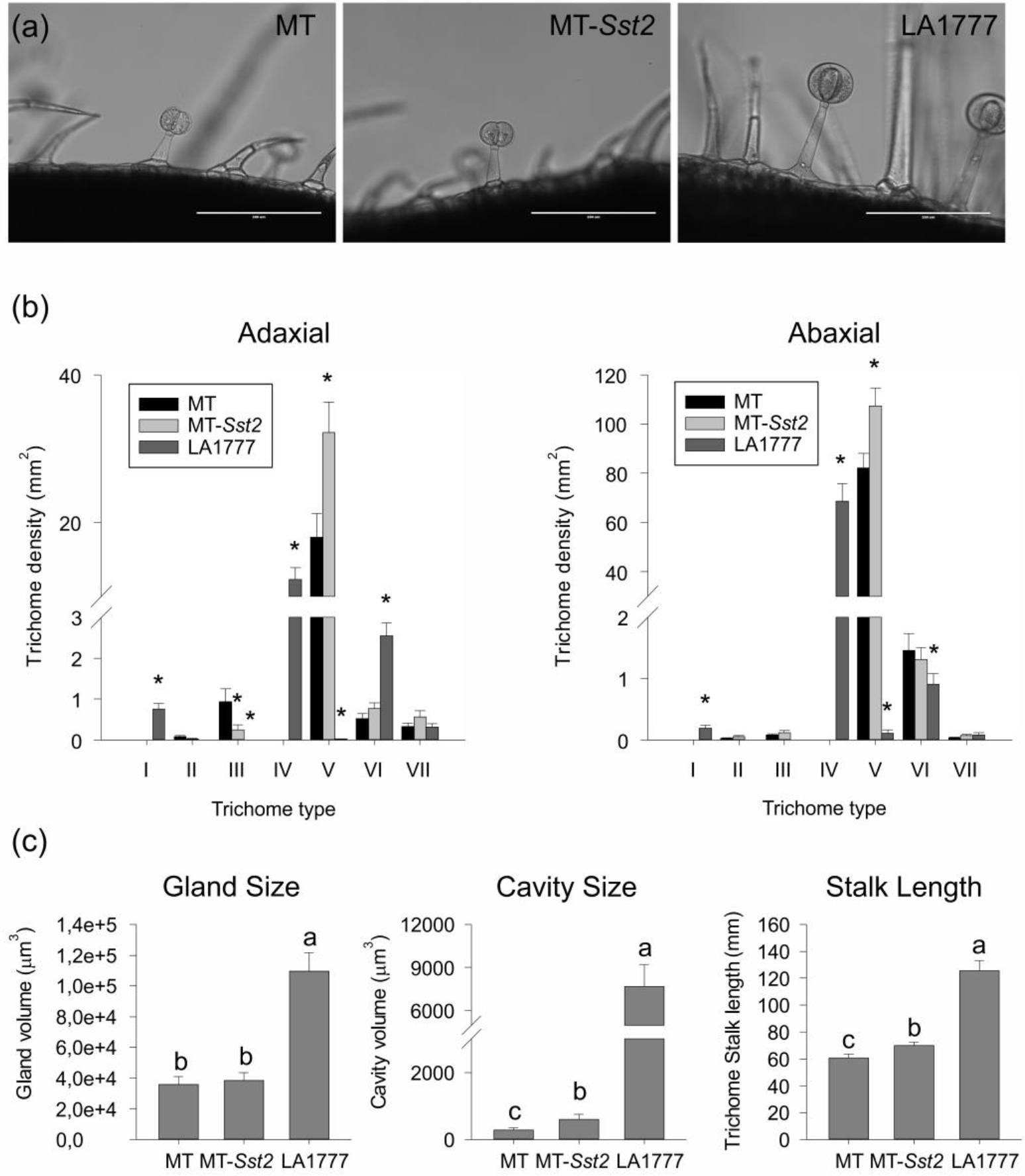
**(a)** Bright field microscopy of trichomes on the leaf surface of representative 45-days old plants of Micro-Tom (MT), MT-*Sst2*, and *Solanum habrochaites* LA1777. Scale bar = 200 μm. **(b)** Density (mm^2^) of trichome types on adaxial and abaxial leaf surfaces. Data are mean (n=40) for each surface. Asterisks indicate mean significantly different from the control MT, according to Student t-test (P≤ 0.05). **(c)** Trichome gland size, cavity volume and stalk length of type-VI trichomes. Data are mean (±SE) of 20 trichomes of 2 replicate leaves of five plants. Bars indicated with different letters were significantly different according to Fisher’s LSD test (P≤ 0.05) after ANOVA.

We also observed that *S. habrochaites* LA1777 has a high density of type-IV trichomes (Fig. 5B). Type-IV trichomes were previously associated with increased production of acylsugars providing resistance to insect pests in wild species (Simmons and Gurr, 2005). Type-IV trichome are however absent on adult leaves of MT and MT-*Sst2*, indicating that the introgressed segment on chromosome 8 from *S. habrochaites* LA1777 is not involved in controlling the presence of this type of trichomes. Type-V is the most abundant non-glandular trichome found on MT and MT-*Sst2*. Interestingly, we found an increased number of (non-glandular) type-V trichomes on both leaf surfaces in MT-*Sst2*. Approximately 2-fold more type-V trichomes were observed on the adaxial leaf surface of MT-*Sst2* compared with MT background (Fig. 5B).

The majority of glandular trichomes found in MT and MT-*Sst2* was type-VI. We next analysed the gland and internal cavity size of the type-VI glandular trichomes (Fig. 5A and C). There is no significant difference between the MT and MT-*Sst2* gland size. However, the size of the internal cavity is increased in MT-*Sst2* compared to MT. The introgression line also appears to exhibit a (subtle but significant) increase in stalk length compared to MT, but still the stalk is much shorter than that of type-VI trichomes found in LA1777 (Fig. 5A and C).

### Herbivore resistance of the Micro-Tom line with increased amounts of santalene and bergamotene

In order to verify if the increase in plastid-derived sesquiterpenes in type-VI trichomes of MT-*Sst2* would result in improved resistance to herbivores, we conducted no-choice bioassays using four relevant pests in tomato. In the whitefly bioassay, survival of whiteflies did not differ between MT-*Sst2* and MT plants, while the LA1777 displayed a high reduction in the survival of this pest. Almost 80% of the whiteflies survived on MT-*Sst2*, whereas less than 40% survived on the wild tomato species (Fig. 6A).

**Fig. 6.**
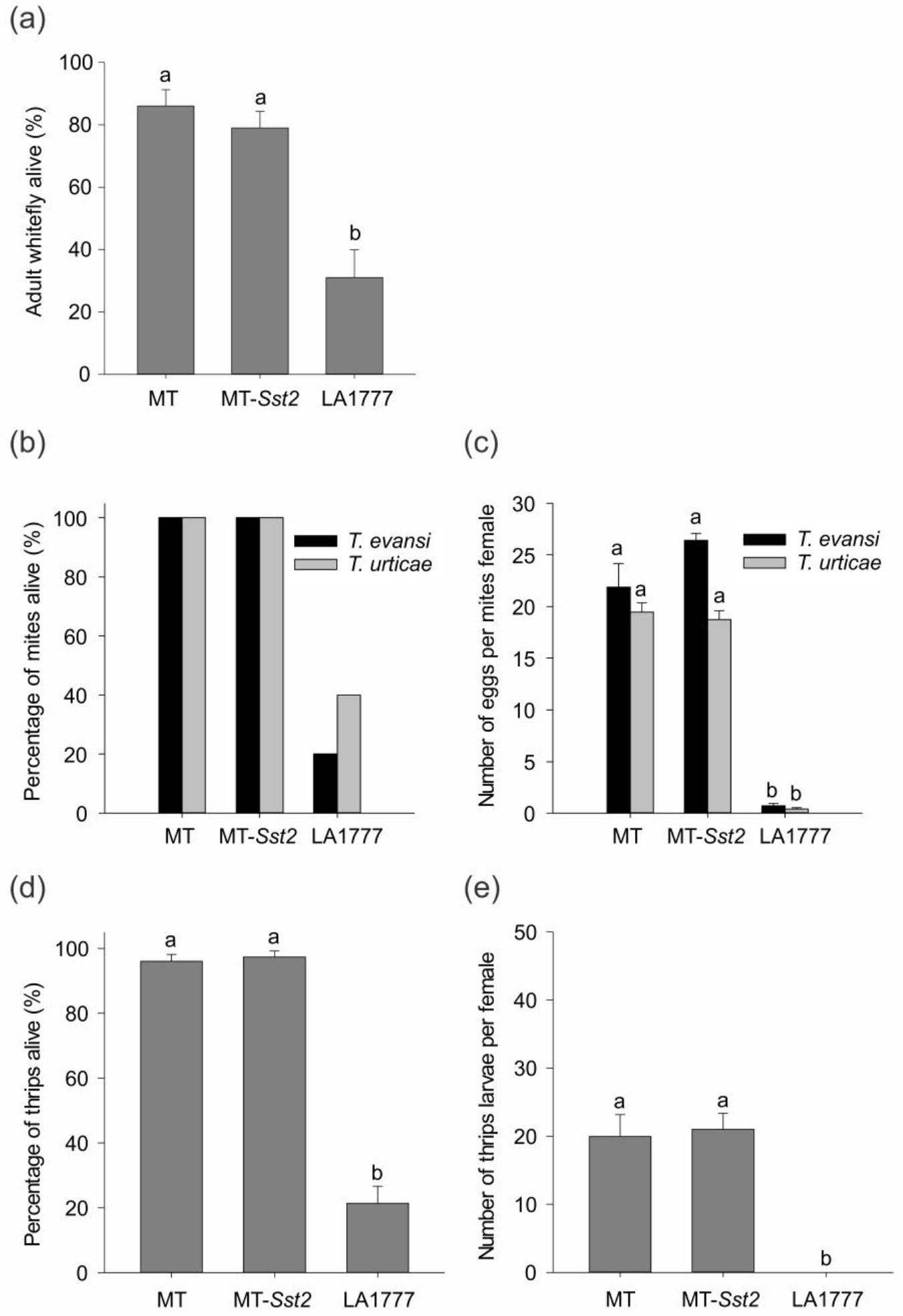
Herbivory tests performed on Micro-Tom (MT), MT-*Sst2* and *Solanum habrochaites* LA1777 genotypes. **(a)** Percentage of adult whitefly *Bemisia tabaci* alive after five days of feeding on leaves. Data are means (±SE) of five plants, each with two cages. Bars indicated with different letters were significantly different according to Fisher’s LSD test (P≤ 0.05) after ANOVA. **(b)** and **(c)** Female spider mite (*Tetranychus evansi* and *Tetranychus urticae)* survival and number of eggs after two days of feeding on plants. Bars indicated with different letters were significantly different according to Fisher’s LSD test (P≤ 0.05) after ANOVA. **(d)** and **(e)** Percentage of adult thrips alive and number of thrips larvae per female that emerged after two days on leaf discs. Bars indicated with different letters were significantly different according to Fisher’s LSD test (P≤ 0.05) after ANOVA.

Next, bioassays using the defense-suppressing spider mite *Tetranychus evansi* and the defense-inducing spider mite *Tetranychus urticae* were performed. Both MT and MT-*Sst2* showed 100% of both spider mites species surviving, while on *S. habrochaites* LA1777 only approximately 20% of *T. evansi* and 40% of *T. urticae* survived after 2 days (Fig. 6B). Oviposition rates of both spider mite species were equal on MT-*Sst2* and MT leaf disc, as they were strongly reduced in the wild species (Fig. 6C).

Finally, we conducted a bioassay with the western flower thrips (*Frankliniella occidentalis*) comparing adult survival on MT, MT-*Sst2* and the wild species. Again, there were no significant differences in survival or egg hatching observed as a result of the *Sst2* introgression (Fig. 6D and E). Female thrips were still able to oviposit in both genotypes and the larvae hatched from the eggs reaching the larval stage. Conversely, significant reductions in the number of surviving adults and the number of emerged larvae were observed for the wild species.

## DISCUSSION

### *Solanum habrochaites* plastid-derived sesquiterpene synthesis can be transferred to type-VI trichomes of cultivated tomato

We successfully introgressed the *SsT2* gene cluster responsible for the biosynthesis of plastid-derived sesquiterpene pathway from *S. habrochaites* LA1777 into the genetic model system cv. Micro-Tom (MT) (Fig 1A). The genes transferred were expressed into the tomato type-VI trichomes (Fig. 2) and resulted in the production of high levels of plastid-derived sesquiterpenes (Fig. 3C).

The allelic dosage influenced the plastid-derived sesquiterpenes levels produced in type-VI trichomes. The levels of all plastid-derived sesquiterpenes were higher in the homozygous *Sst2*/*Sst2* plants than the hemizygous *sst2/Sst2* (Fig.4). This can be explained by the presence of just one copy of both *zFPS* and *SBS* alleles in hemizygous *sst2/Sst2* plants. It can also be the result of the presence of both *SlNDPS1* and *zFPS* in the hemizygous *sst2/Sst2* acting in the same compartments (plastids) and competing for a limited amount of IPP and DMAPP in type-VI trichomes (Dudareva *et al.*, 2005; Besser *et al.*, 2009; Schilmiller *et al.*, 2009). However, it is unlikely that plastid-derived monoterpene production can limit the substrate for the production of sesquiterpenes in plastids, since the levels of monoterpenes are much lower than the plastid-derived sesquiterpenes (Fig. S2, Fig. 4) (Banerjee *et al.*, 2013). In addition, due to differences in substrate affinity, zFPS would use the IPP or DMAPP to produce *Z,Z*-FPP more efficiently than SlNDPS1 uses to produce NPP in the plastids (Sallaud *et al.*, 2009; Kang *et al.*, 2014).

Despite the fact that the introgression line MT-*Sst2* produced significantly higher levels of santalene and bergamotene in type-VI glands compared to LA1777 (Fig 3), we did not detect sesquiterpene derivatives in MT-*Sst2* as they are present in the wild species (Fig.3A and C). The absence of *α*-santalenoic and *α*- and *β*-bergamotenoic acids in the MT-*Sst2* line likely explains the higher levels of santalene and bergamotene, compared to the wild species. In *S. habrochaites* LA1777, santalene and bergamotene are converted to derivatives identified as sesquiterpene carboxylic acids, *α*-santalenoic, and *α*- and *β*-bergamotenoic acids (Coates *et al.*, 1988). Notably, high sesquiterpene levels in MT-S*st2* compared to LA1777 provide an indirect evidence that santalene and bergamotene are used as precursors for further metabolism into corresponding alcohol and acids derivatives in LA1777 (Frelichowski and Juvik, 2005; Besser *et al.*, 2009; Gonzales-Vigil *et al.*, 2012).

The relative transcript levels of *zFPS* and *SBS* were higher in LA1777 compared to the introgressed line (Fig. 2). Since MT-*Sst2* and LA1777 share the same chromosomal segment comprising the *SsT2* locus, it was expected that they have the same *cis*-regulatory elements controlling the expression of the *zFPS* and *SBS* genes. The lower expression of these genes in MT-*Sst2* may suggest the involvement of a set of yet unknown trans-regulatory elements (e.g. transcription factors and other regulatory elements in different chromosomal regions) necessary to increase the expression of the genes present in the *SsT2* locus. Therefore, our results point at a role for additional genetic components that have not been introgressed. It has been shown that poor terpene producing genotypes have also drastically reduced transcript levels for key steps in the terpene biosynthesis pathway (Tissier, 2012), suggesting that the transfer of additional components enhancing *zFPS* and *SBS* expression could increase the content of sesquiterpenes in MT-*Sst2* even further. Up to now, only a few transcription factors were proven to be involved in the regulation of terpene pathways in tomato (Spyropoulou *et al.*, 2014; J., Xu *et al.*, 2018), though these were not implicated as positive regulators of *zFPS* or *SBS* specifically.

We also noted that the concentration of the cytosolic-derived sesquiterpene α-humulene was higher in homozygous *Sst2/Sst2* plants when compared to homozygous *sst2/sst2* and hemizygous *sst2/Sst2* plants. The TPS12 responsible for the production of β-caryophyllene and α-humulene is encoded by a gene in the *SsT1* locus on chromosome 6, which is clearly outside of the region introgressed. Therefore, the *TPS12* in the *SsT1* locus is likely to have the same alleles for the three genotypes and it could not be the cause of the high amount of α-humulene in the homozygous *Sst2/Sst2* plants. Since α-humulene coelutes with β-bergamotene (Hoeven van der *et al.*, 2000), we cannot exclude the possibility that the level of α-humulene in MT-*Sst2* is overestimated.

### Introgression of Sst2 appears to affect type-VI trichome morphology

The MT-*Sst2* line with augmented contents of sesquiterpenes in type-VI trichomes displayed an increased glandular cavity volume (Fig 5C). A possible explanation for this is that the boost in the total amount of terpenes could result in a physical pressure in the cavity wall, forcing the internal cavity to inflate like a balloon, as similarly suggested by Ben-Israel et al., (2009). However, the subtle increase in the internal cavity of MT-*Sst2* was not paired with an altered external gland shape. Modification in external gland shape depends both on genes related to cell wall remodelling (Bennewitz *et al.*, 2018) and on synthesis and accumulation of very high levels of compounds into the gland. Thus, a combination of genes controlling the high flux of metabolites with genes controlling the cell wall remodeling might push the gland to expand, creating the characteristic round type-VI trichome of *S. habrochaites*.

Stalk length is one of the morphologic characteristics used to identify the different types of trichomes in *Solanum* (Simmons and Gurr, 2005). The slightly increased type-VI trichome stalk length observed in MT-*Sst2* (Fig. 5C) is unlikely the result of the effect of the biosynthetic enzymes encoded on the *SsT2* locus. Recently Xu et al., (2018c), demonstrated that the downregulation of *SlMYC1*, a bHLH transcript factor, results in plants with shorter type-VI trichome stalks. Since *SlMYC1* (*Solyc08g005050*) is also located on top of chromosome 8, we specifically searched for this transcript factor in our introgressed line. Hence, the presence of the wild *ShMYC1* allele inside the region introgressed was confirmed by CAPS marks (Fig 1B and S3). In general, *S. habrochaites* species exhibit a longer type-VI trichome stalk compared to cultivated tomatoes (Simmons and Gurr, 2005; Bergau *et al.*, 2015). The biological role for a higher stalk, or a taller trichome in the wild species is not described, but it is tempting to speculate that though the effect is marginal, the altered stalk length could be the result of replacement of *SlMYC1* with *ShMYC1*.

Furthermore, MT-*Sst2* displayed an increased number of type-V non-glandular trichomes compared to both MT and LA1777 (Fig 5B). It was shown earlier that there appears to be a negative correlation between densities of glandular type-IV trichomes and the non-glandular type-V trichomes (Vendemiatti *et al.*, 2017). The lower density of type-V trichomes in LA1777 could therefore be related to its high density of type-IV trichomes. The adult leaves of MT-Sst2 and MT do not have type-IV glandular trichomes (Fig. 5B). This lack of type-IV trichomes in cultivated tomato has been linked to the transition from the juvenility to the adult phase (Vendemiatti *et al.*, 2017). So, the region introgressed in MT-*Sst2* is most likely not involved in heterochrony and/or type-IV trichome development, though we cannot discard the possibility that the region introgressed harbors genes controlling the density of type-V trichomes, as we cannot explain the increased type-V density in the introgression line compared to MT.

### Genetic factors outside the *SsT2* metabolic cluster are required to produce sesquiterpene carboxylic acids (SCA) terpenoids and insect resistance

Even though MT-*Sst2* produced relatively high levels of santalene and bergamotene, compared to *S. habrochaites* LA1777, this did not confer resistance to any of the herbivores tested here (Fig. 6). Herbivore resistance observed in the wild species is therefore likely due to the sesquiterpene carboxylic acids (SCAs) derivatives absent in MT-*Sst2*. We cannot rule out the contribution of type-IV trichomes-derived acylsugars in LA1777 (Kim *et al.*, 2012), but it has been shown before that the presence of SCAs in LA1777 modulates larval feeding behavior and survival of two tomato insect pests (Frelichowski and Juvik, 2001). On the other hand, transgenic *S. lycopersicum* producing the *S. habrochaites* sesquiterpene 7-epizingiberene, product of the expression of *zFPS* and *ShZIS*, probably an allelic variant of *ShSBS*, did display a clear toxicity phenotype against spider mites (Bleeker *et al.*, 2012). Transference of the complete SCA route to MT-*Sst2* will answer this question on the contribution of SCA in herbivore resistance.

Up to now, little is known about the enzymes catalyzing the formation of SCAs from sesquiterpenes in LA1777. It can be hypothesized that cytochrome P450 enzymes (which often hydroxylate terpenes) play a role. In *Santalum album* (Santalaceae) santalenes (α-, β- and *epi*-β-santalene) and α-*exo*-bergamotene are further metabolized into sesquiterpene alcohols α-, β-, and *epi*-β-santalol and α-*exo-*bergamotol by a CYP76F cytochrome P450 (Diaz-Chavez *et al.*, 2013). Nevertheless, co-expression of *S. album* santalene/bergamotene oxidase *SaCYP76F39v1* in transgenic tobacco plants did not hydroxylate santalene or bergamotene (Yin and Wong, 2019). In *Tanacetum cinerariifolium (*Asteraceae), two oxidation reactions convert trans-chrysanthemol into trans-chrysanthemic acid (H., Xu, Moghe, *et al.*, 2018). Using tomato transgenic lines, Xu *et al.* (2018a) also showed that expressing chrysanthemyl diphosphate synthase from *Tanacetum cinerariifolium* together with two an alcohol dehydrogenase and aldehyde dehydrogenase from the *S. habrochaites* LA1777 were sufficient for the transgenic fruits to produce trans-chrysanthemic acid.

Although the introgressed region in MT-*Sst2* contains both an alcohol dehydrogenase (ADH) and P450s from *S. habrochaites* LA1777, it seems that the genes required for the conversion of the terpenes to derivatization of the sesquiterpenes to their alcohols or carboxylic acids lie outside the metabolic cluster on chromosome 8. Peripheral pathway genes located outside the core metabolic cluster in different chromosomes have been described before for tomato and other species (Nützmann *et al.*, 2016). The fact that we did not detect any sesquiterpene (intermediate) derivative in MT-*Sst2* indicates that the alcohol dehydrogenases and cytochrome P450s introgressed must have different functions.

Altogether, this study demonstrates that the plastid-derived sesquiterpene synthesis from *S. habrochaites* can be transferred to type-IV trichomes of cultivated tomato. However, additional genetic components from the wild species should be transferred to acquire herbivore resistance in cultivated tomato. The results presented here indicate that at least part of these genetic components is likely to encode: i) enzymes that convert santalene and bergamotene into their carboxylic acids, ii) transcription factors modulating the expression *zFPS* and *SBS* and iii) enzymes and possible regulatory genes for cell wall remodeling and enlargement of the internal cavity of the trichome gland. The introgressed line presented here, in the model system Micro-Tom, can provide for rapid introgression and transgenic manipulation of the additional genetic components involved in sesquiterpene metabolism and type-VI trichome morphology.

## MATERIAL AND METHODS

### Plant Material

Seeds from Micro-Tom (MT) were donated by Dr. Avram Levy (Weizmann Institute of Science, Israel) in 1998 and maintained through self-pollination as a true-to-type cultivar since then. The *lutescent 1* mutation was introgressed into MT from its original background as described previously in Carvalho *et al.* (2011). Seeds from *Solanum habrochaites* LA1777 were obtained from the Tomato Genetics Resource Center (TGRC - University of California).

The sesquiterpene synthase 2 pathway from *S. habrochaites* LA1777 was introgressed into MT background by allelic substitution making use of the morphological marker MT-*lutescent 1*, which maps on the same arm of chromosome 8 (https://tgrc.ucdavis.edu/) (Fig.1). Briefly, pollen from *S. habrochaites* LA1777 were collected and used to fertilize emasculated MT-*lutescent 1* flowers. The F1 obtained was used as pollen donor for MT-*lutescent 1* plants and this procedure was repeated in the successive backcrossing (BCs). In each BC, we screened for reduced plant size (MT-like phenotype) and the absence of the *lutescent 1* phenotype (Fig. S1), which is an indicative of the presence of the LA1777 genes in the *SsT2* locus. After self-pollination in BC_6_F_2_ generation, we screened plants for the presence of the same sesquiterpenes compounds found in the wild parental species. The resulting homozygous MT-*Sst2* genotype was considered a near-isogenic line (NIL).

Plants were grown in a greenhouse with 30/26°C temperature day/night and 60–75% ambient relative humidity, 11.5 h/13 h (winter/summer) photoperiod, sunlight 250–350 μmol photons m^−2^ s^−1^ PAR irradiance. Seeds were germinated in bulk in 350 mL pots with a 1:1 mixture of commercial potting mix Basaplant^®^ and expanded vermiculite, and was supplemented with 1 g L^−1^ 10:10:10 NPK and 4 g L^−1^ dolomite limestone (MgCO_3_ + CaCO_3_). Upon the appearance of the first true leaf, seedlings of each genotype were individually transplanted to 150 mL pots containing the soil mix described above, except that NPK supplementation was increased to 8 g L^−1^.

### Genetic and Physical Mapping of the introgressed *Sst2* genes

Genomic DNA isolation was extracted from leaflets using the method described by Fulton, Chunwongse & Tanksley (1995) with minor modifications. Molecular mapping using Cleaved Amplified Polymorphic Sequence (CAPS) markers was performed as previously described by Shavrukov (2016). Details of tomato genetic maps and chromosome 8 molecular markers can be accessed through the Solanaceous Genomics Network (http://solgenomics.net/). Primers and restriction enzymes yielding CAPS between tomato and *S. habrochaites* LA1777 are detailed in the Supplemental Table S1.

### Trichome counts and phenotyping

Counting of trichomes density (mm^2^) were performed on leaflets taken from mature fifth leaves (counting from cotyledons) according to the methodology described by Vendemiatti *et al.* (2017). Both leaf surfaces were dissected along the longitudinal axis in 15 × 3 mm strips covering the middle section of the leaf blade (avoiding the primary veins). The strips were fixed on microscope slides using transparent nail polish. Five individuals per genotype were sampled, and four different strips were analyzed per plant. Images were taken using a Leica S8AP0 stereomicroscope (Wetzlar, Germany) magnifying glass set to 50× magnification, coupled to a Leica DFC295 camera (Wetzlar, Germany).

The morphology of type-VI trichomes was examined under an EVOSfl (www.thermofisher.com) inverted microscope. Lateral leaflets strips were submerged in water under microscope slides and images of type-VI trichomes were taken. All trichome measurements were performed on images of 5 plants per genotype using ImageJ software version 1.4.1. Gland volume and cavity volume were calculated using the volume of the prolate ellipsoid formula: *V* = *4*/*3* × *π* × *a* (vertical axis) × *b*^*2*^ (horizontal axis).

### GC-MS quantification

For GC-MS volatile terpene quantification 300 individual type-VI trichome glands were collected from leaves in adult vegetative phase (fifth leaf from the cotyledons) with a glass pulled Pasteur pipette under a Leica MZFLIII microscope (www.leica-microsystems.com). The terpene extraction was conducted according to the methodology described by Xu *et al.* (2018c). The collected glands were dissolved in 150 μL of hexane plus 0.5 ng μL of benzyl acetate (Sigma-Aldrich; www.sigmaaldrich.com) as an internal standard. Na_2_CO_3_ (Sigma-Aldrich) was used to remove water from the hexane. Volatiles were separated using an Agilent (www.agilent.com) 7890A gas chromatograph, attached to an Agilent 7200 accurate-mass quadrupole time-of-flight mass spectrometer. Here, 2μL of the sample was injected heated to 275°C in the injector port and separated on an HP-5ms column (0.25 mm in diameter, 30 m in length, with 0.25 μm film thickness) using Helium as carrier gas (flow rate 1 mL/min). The oven temperature was maintained at 40°C for 3 min and increased by 15°C per min until it reached 250°C and maintained for 3 min. Identification of the compounds was based on the retention time of the chromatographic peaks and their corresponding mass spectra, which were compared to terpene standards and data libraries. Quantification of peak areas was performed using Masshunter Qualitative Analysis software (Agilent). Peak areas were corrected for the internal standard and quantified using the available terpene standards. For compounds without terpene standards, we used the β-caryophyllene standard as a reference. Terpene concentration was calculated per trichome gland (ng/gland) using the peak areas relative to the internal (benzyl acetate) and terpene standards available.

### Whitefly bioassay

*Bemisia tabaci* (former biotype B; Middle East Asia Minor I-II (MEAM)) population was maintained in a climatized chamber (Snijders Tilburg; T 28°C, 16-h light, RH 75%) on cucumber plants prior to the experiment. For no-choice assay twenty adult whiteflies were randomly taken from the population, anesthetized with CO_2_ and placed in a clipcage (2.5 cm diameter; Bioquip). Two clipcages were attached in two different leaflets per plant. Five plants per genotype were used. The plants were kept inside of a closed greenhouse compartment (28 °C, RH 65%) and after 5 days, the number of whiteflies alive was recorded.

### Spider Mite bioassay

A non-choice performance assay was set-up using two species of spider mites. The two spotted spider mite *T. urticae* Koch Viçosa-1 and the red spider mite *T. evansi* Baker & Pritchard Viçosa-1 were initially collected from infested tomato plants (Sarmento *et al.*, 2011). Before the experiments, *T. urticae* mites were maintained on detached leaves of *S. lycopersicum* cv. Santa Clara and *T. evansi* mites were maintained on detached leaves of *S. lycopersicum* cv. Castlemart following standard procedures (Ataide *et al.*, 2016). The rearings were maintained in a climate room at 25 °C, a 16/8 h light regime with 300 μE m^−2^ s^−1^, and 60% RH.

For each plant genotype, 15 leaf discs of 15 mm were made from the fifth leaf (counting from the cotyledons). The leaf discs were placed on 1.5% daishin agar (Duchefa Biochemie bv, Haarlem, The Netherlands) that was poured in small cups (3 cm diameter × 2 cm height) with their adaxial side facing up. On each leaf disk, a single 2-day-old adult female of *T. urticae* or *T. evansi* was placed using a soft paintbrush. Mites were confined into each cup and ventilation was assured by a 1 cm^2^ opening on the lid that was covered with mite-proof mesh (pore size of 80 μm). The cups were maintained in a climate room at 25°C, a 16/8 h light regime with 300 μE m^−2^ s^−1^, and 60% RH. Two days after infestation, spider mite survival was recorded and the average fecundity (number of eggs laid per female) was calculated using those spider mites that were alive after the two-day period for the calculations.

### Thrips bioassay

A non-choice performance thrips bioassay was set up using the western flower thrips *Frankliniella occidentalis* (Pergande). The thrips colony was kept in the laboratory inside cages where bean pods were provided and supplemented with pollen as previously described by Muñoz-Cárdenas et al., (2017). For each plant genotype, 15 leaf discs of 15 mm diameter were made. Similar to the spider mites set-up, the experimental arena consisted in cups (3 cm diameter × 2 cm height) filled with 1.5% Daishin agar on which one leaflet was placed with the adaxial side up. Five adult females were collected from the colony with the help of a 1 ml pipette tip attached to a vacuum and were released inside each cup through a small opening on the side of the cup, that was otherwise sealed with parafilm. The cups were maintained in a climate room at 25°C, a 16/8 h light regime with 300 μE m^−2^ s^−1^, and 60% RH. 72 hours after the release of the females, the adult thrips were removed and their survival (number of alive thrips) was scored with the help of a dissecting stereoscope. The number of larvae that emerged from the eggs laid during the experiment was assessed 7 days after the beginning of the experiment.

### RNA Isolation and Quantitative RT-PCR

Total RNA was extracted from trichomes isolated by shaking stems in liquid nitrogen with a vortex mixer. Total RNA was isolated using Trizol reagent (Invitrogen) according to the manufacturer’s instructions. RNA treated with TURBO DNase (Ambion; www.thermofisher.com) were reverse-transcribed to generate first-strand cDNA using RevertAid H Minus Reverse Transcriptase (Fermentas; www.thermofisher.com). cDNA was used as a template for quantitative RT-PCR (qRT-PCR). PCR reactions were performed using HOT FIREPol EvaGreen qPCR Mix Plus (Solis Biodyne; www.sbd.ee) and analyzed in an ABI 7500 Real-Time PCR System (Applied Biosystems; www.appliedbiosystems.com). Two technical replicates were analyzed for at less three biological samples, together with template free reactions as negative controls. Transcript abundances were normalized to *Rubisco conjugating enzyme 1* (*RCE1*) expression. Detailed primers information is described in the Supplemental Table S2.

### Experimental Design and Statistical Analysis

Statistical analyses were done using SigmaPlot 11.0 for Windows. The experiments were arranged in a completely randomized design. All data were tested for normality and equal variance by Kolmogorov-Smirnov tests. The means were further analyzed by two-tailed Student’s *t*-test (P≤ 0.05) or Fisher’s LSD test (P≤ 0.05) after one-way ANOVA in multiple comparisons. For data that do not assume a specified variance or normality, we performed ranking tests Wilcoxon rank sum for pairwise comparisons and Kruskal–Wallis one-way analysis for multiple groups.

## ACKNOWLEDGMENTS

The authors thank L.E.P.P., P.B., R.S. and S.M.A.’s lab members for laboratory and greenhouse assistance.

---

**Fig. S1.**
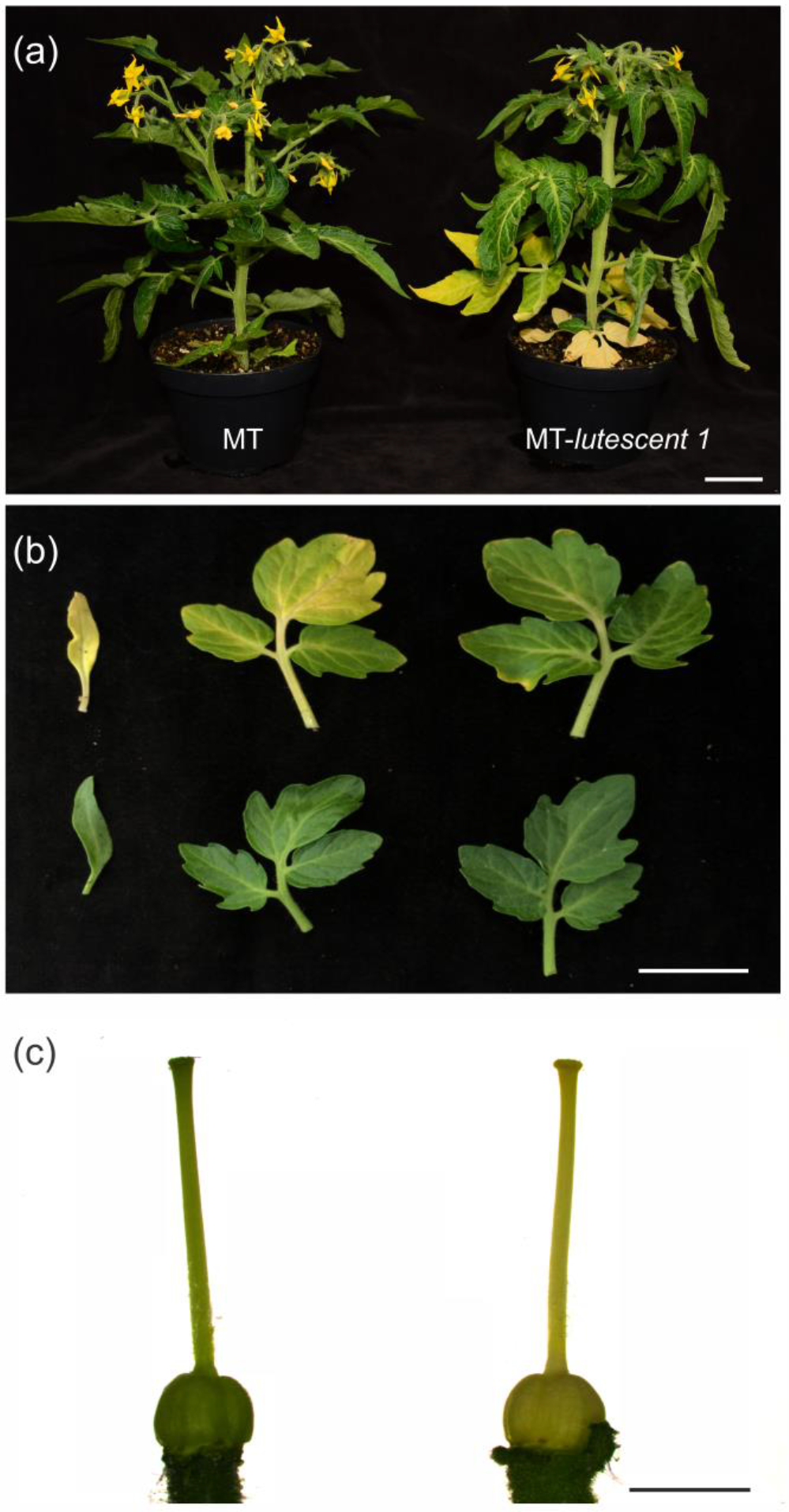
**(a)** Representative MT and the MT near isogenic line (NIL) harboring the *lutescent 1* (l/l) mutation (MT-*lutescent 1*) which shows a premature and progressive chlorophyll loss that was used as a morphological marker for genetic introgression. Bar = 2 cm. **(b)** MT-*lutescent 1* leaf phenotype (top) displaying premature and progressive chlorophyll loss on cotyledons and the first and second leaves, compared with MT (bottom), which do not present senescent-like leaves at this developmental stage. Bar = 3 cm. **(c)** MT-*lutescent 1* pistil (right) with chlorophyll loss, compared with MT pistil (left). Bar = 2 mm. The photos were taken from 45-days old plants.

**Fig. S2.**
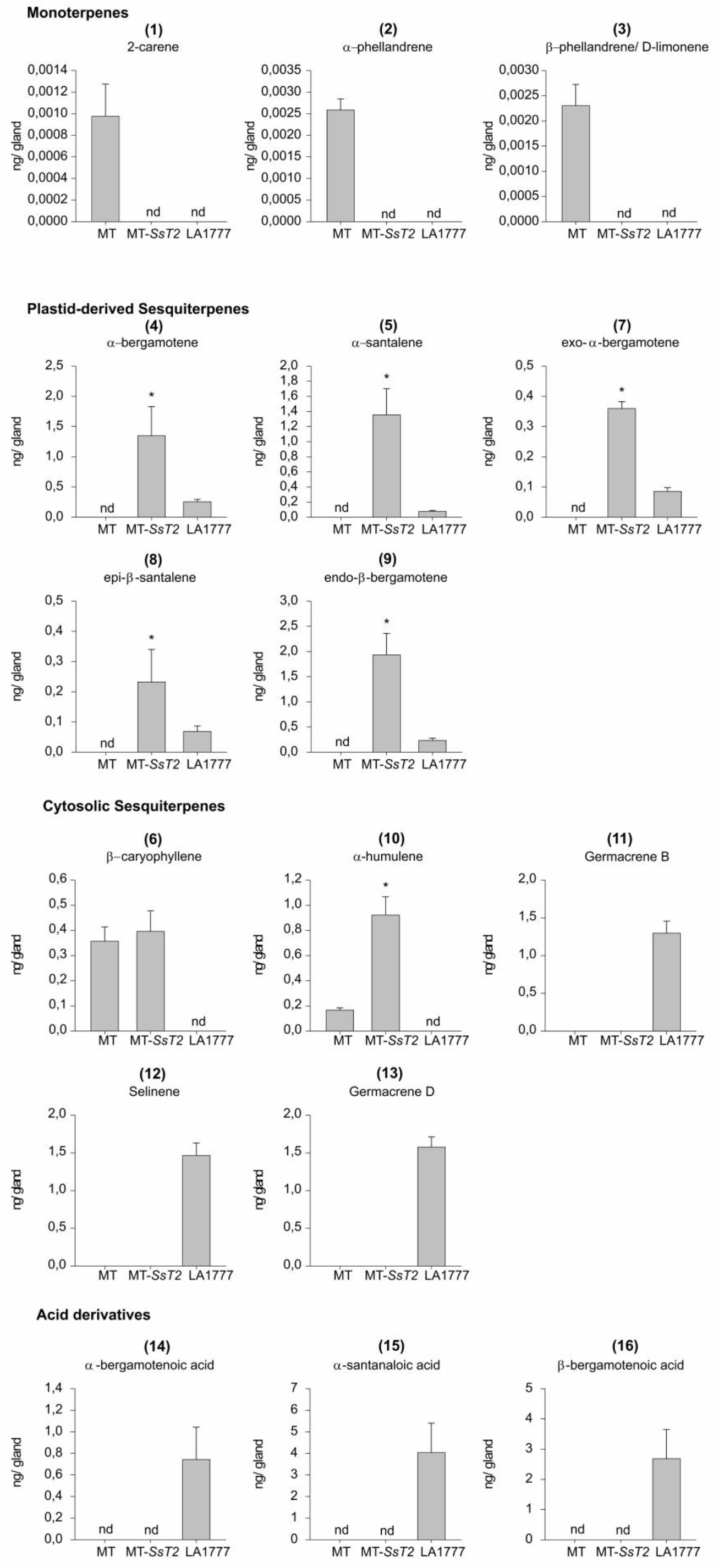
Volatile terpene levels in type-VI glandular trichomes from Micro-Tom (MT), MT-*Sst2* and the wild species *Solanum habrochaites* LA1777. The data show the amount of each compounds present in type-VI glandular trichomes. Each data point represents the mean + SE of five biological replicates. For each sample, 300 type-VI glandular trichomes were collected with a glass capillary before GC-MS. Means indicated with an asterisk were significantly different according to t-test (P≤ 0.05). nd, Not detected.

**Table S1.**
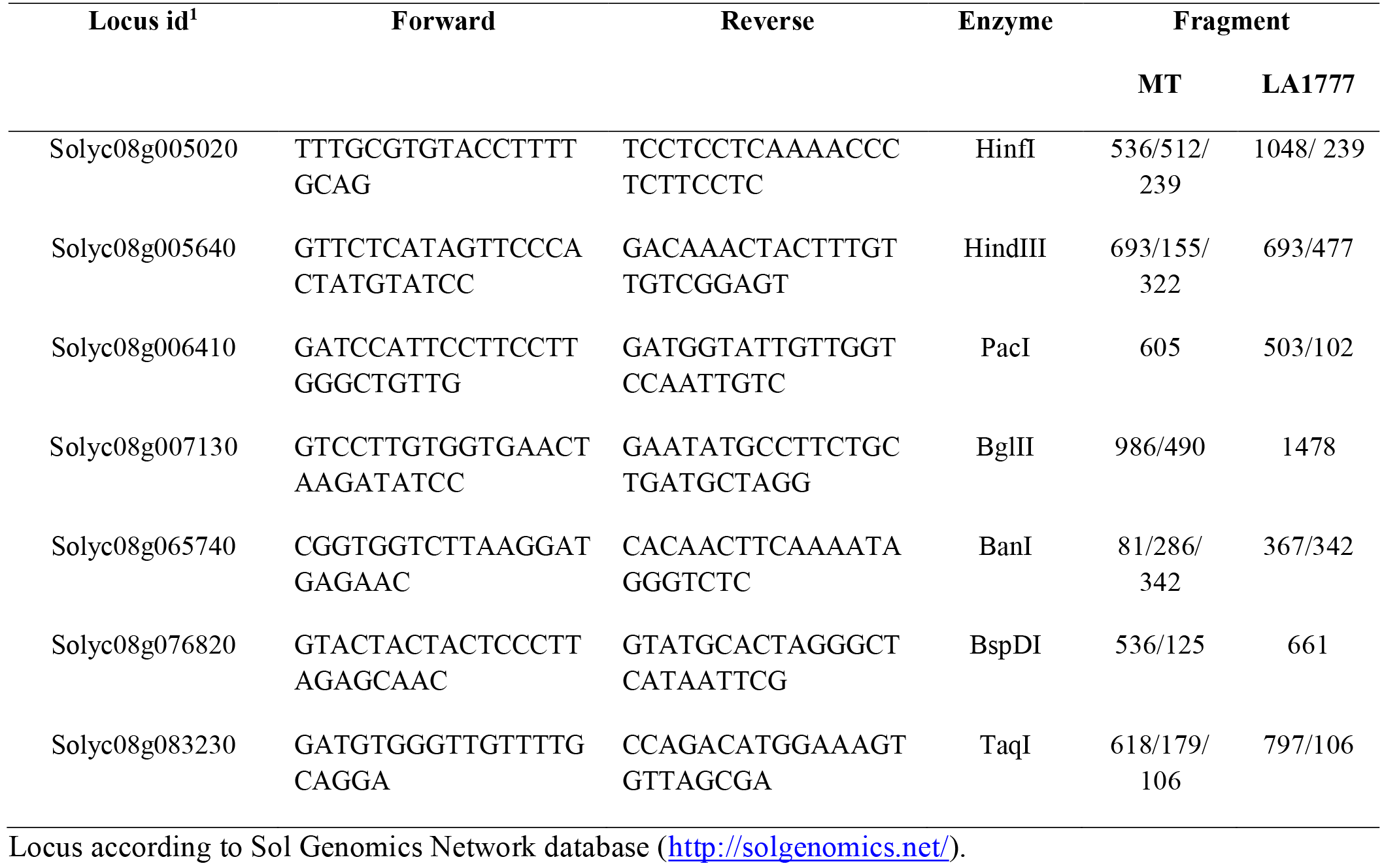
Oligonucleotide sequence used for CAPS markers.

**Fig. S3.**
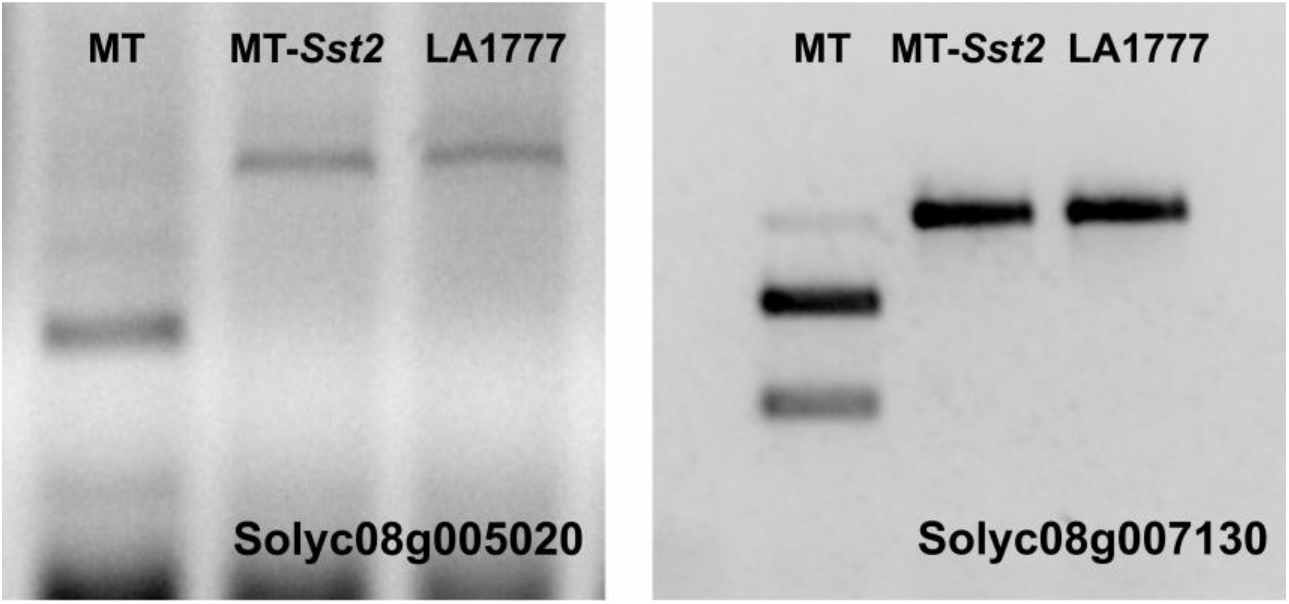
Electrophoresis gels showing the positive genetic markers used to differentiate MT-*Sst2* from MT.

**Table S2.**
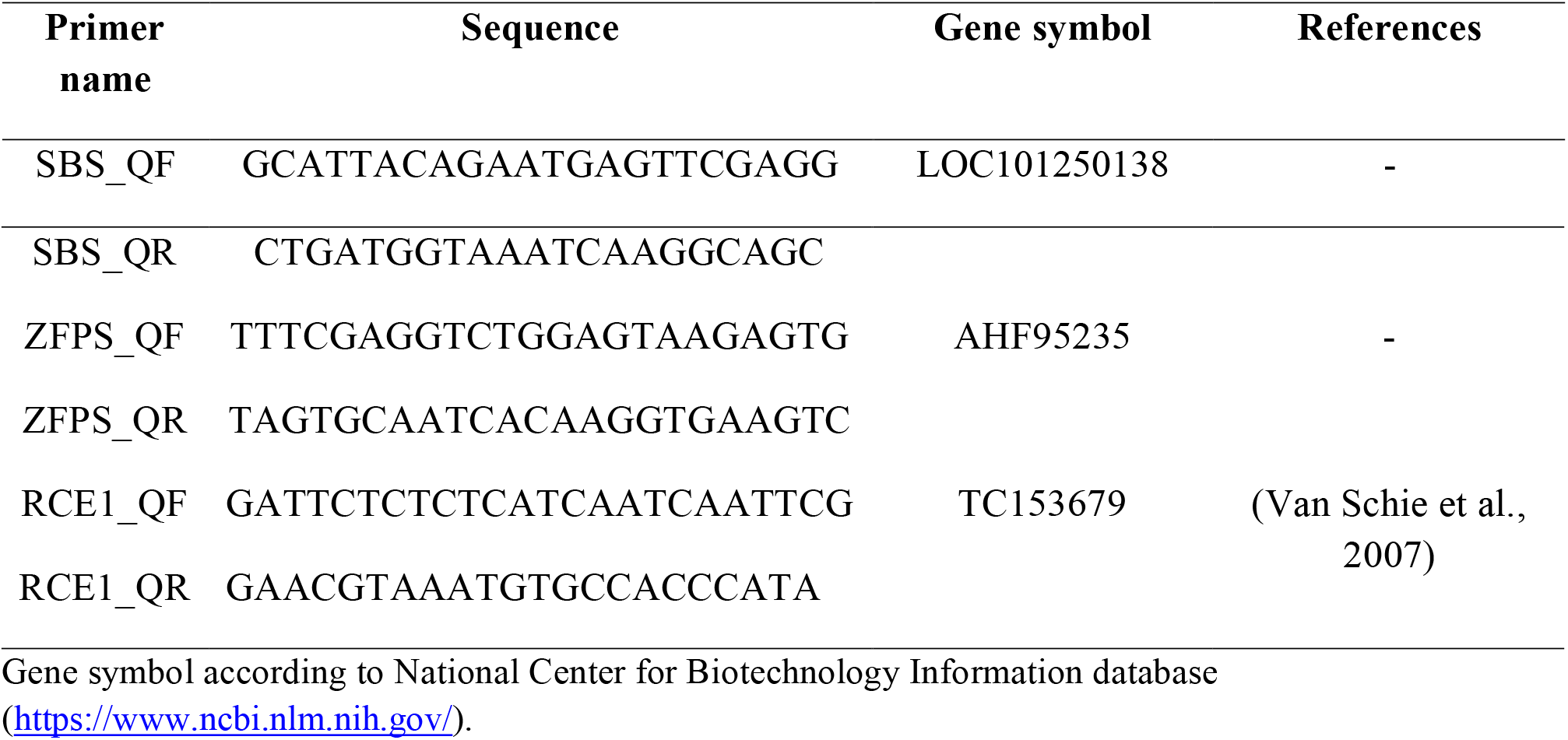
Oligonucleotide sequence used for quantitative PCR analyses.

